# The complex development of psoralen-interstrand crosslink resistance in *Escherichia coli* requires AcrR inactivation, retention of a *marbox* sequence, and one of three MarA, SoxS, or Rob global regulators

**DOI:** 10.1101/2024.12.03.626702

**Authors:** Travis K Worley, Ayah H Asal, Lo Cooper, Charmain T Courcelle, Justin Courcelle

## Abstract

Crosslinking agents, such as psoralen and UVA radiation, can be effectively used as antimicrobials and for treating several dysplastic conditions in humans, including some cancers. Yet, both cancer cells and bacteria can become resistant to these compounds, making it important to understand how resistance develops. Recently, several mutants were isolated that developed high-levels of resistance to these compounds through upregulation of components of the AcrAB-TolC-efflux pump. Here, we characterized these mutants and found that resistance specifically requires inactivating mutations of the *acrR* transcriptional repressor which also retain the *marbox* sequence found within this coding region. In addition, the presence of any one of three global regulators, MarA, SoxS, or Rob, is necessary and sufficient to bind to the *marbox* sequence and activate resistance. Notably, although psoralen is a substrate for the efflux pump, these regulators are not naturally responsive to this stress as neither psoralen, UVA, nor crosslink induction upregulates *acrAB* expression in the absence of mutation.

**Highlights:**

- Psoralen crosslink resistance requires AcrR inactivation and MarA/SoxS/Rob activation
- Psoralen crosslink resistance is mediated by upregulating the AcrAB-TolC efflux pump
- AcrAB-TolC can utilize psoralen as a substrate but not upregulated by this stress
- Acquiring resistance to DNA interstrand crosslinks requires mutation

## 1. Introduction

Psoralen in the presence of UVA irradiation forms DNA interstrand crosslinks and is used in the treatment of psoriasis and vitiligo, as well as in the treatment of cutaneous T-cell lymphoma [1, 2]. The potency of this treatment and similar therapeutics is attributed to its ability to form lethal lesions known as DNA interstrand crosslinks [3–6]. In *E. coli*, a single DNA interstrand crosslink in the genome is sufficient to inactivate the cell [7, 8]. However, the use of psoralen-UVA and other crosslinking agents as antimicrobials and chemotherapeutics can be compromised by the emergence of cells resistant to these drugs [9, 10]. In *Escherichia coli*,several highly resistant mutants to psoralen-UVA interstrand crosslinks have been isolated whose resistance is driven by increased expression of the AcrAB-TolC efflux pump, which protects the DNA and effectively prevents these lethal lesions from forming when psoralen is present in the media [11].

AcrAB-TolC belongs to a highly conserved RND (Resistance-Nodulation-Division) efflux pump family found in Gram-negative bacteria [12–16]. The efflux pump consists of a proton-driven transporter AcrB, a periplasmic adapter protein AcrA, and the TolC transmembrane channel [17–20]. AcrAB-TolC is capable of effluxing a wide variety of structurally dissimilar substrates, including many dyes, detergents, and antibiotics [21–25], making it a primary driver of multiple-antibiotic resistance [23].

The highly resistant mutants were each found to have mutations in in the transcriptional regulator AcrR [11]. *acrR* encodes a TetR family transcriptional regulator that is located immediately upstream of *acrAB* and is divergently transcribed [26]. Based on *lacZ*-fusion and gel mobility shift assays, Ma et. al. demonstrated that AcrR functions as a repressor of *acrAB* that releases upon binding a recognized substrate [27]. Consistent with this, some substrates of the efflux pump, such as rhodamine, ethidium bromide, and proflavine, bind to AcrR [28, 29] and this correlates with a loss of DNA binding activity in vitro [30].

Surprisingly however, deletion of *acrR*’s coding region does not increase resistance to psoralen interstrand crosslinks, suggesting a more complex mechanism of regulation than a simple repressor function is involved [11]. We noted that the first 20 nucleotides of *acrR*’s coding region contains a *marbox*-binding sequence for three closely related global stress regulators, MarA, SoxS and Rob [31]. These three regulators share approximately 50% sequence identity [32, 33] and regulate expression of approximately 50 genes, including *acrA* and *acrB*, in response to various environmental stressors and toxins (Fig. 1 and [31–40]. Using a *lacZ*-reporter construct and gel mobility shift assays, several groups demonstrated that protein binding to the *marbox* upstream of *acrAB* correlated with its expression [27, 40]. This led to a general model that these global stress activators drive *acrAB* expression, with AcrR serving as a secondary repressive modulator.

**Figure 1.**
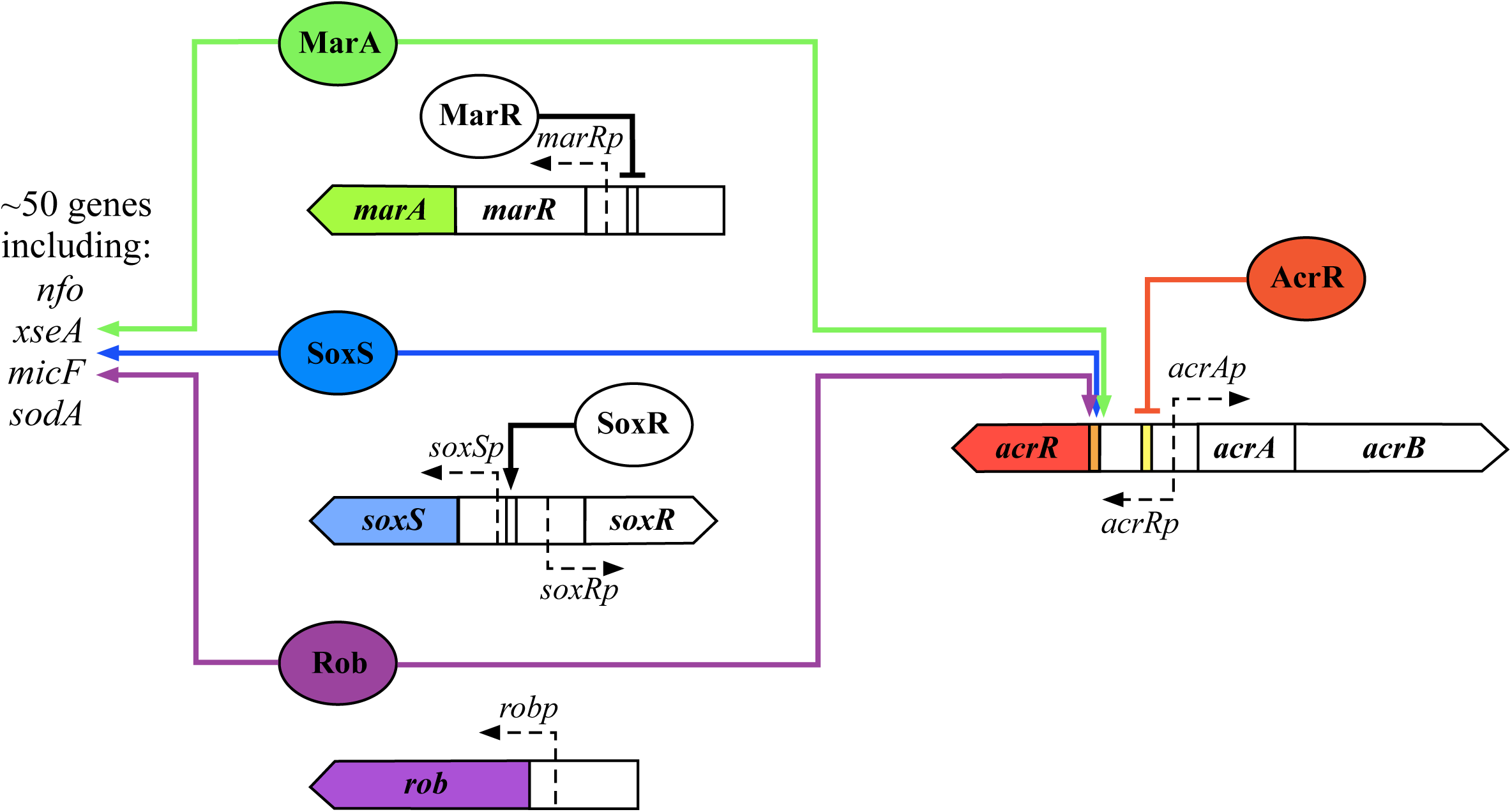
Current model of MarA, SoxS, and Rob global gene regulation. Green, MarA; blue, SoxS; purple, Rob; red, AcrR; yellow, DNA binding sites; orange, Mar/Sox/Rob binding site (*marbox*). Arrows indicate activation, while interruption of the end of a line indicates repression (derived from models and data presented in [26, 31, 33, 36, 37, 45]).

Given the importance of DNA interstrand crosslinks in antimicrobrial and chemotherapeutic therapies, here we sought to characterize the mechanism by which resistance was achieved in these mutants acquired their resistance. We found that although the pump confers resistance to crosslinks, it is not naturally responsive or upregulated in their presence. Resistance relies on mutations that inactivate the AcrR repressor but retain the *marbox* sequence within the gene’s coding region. The resistance can then be activated by the presence of any one of the three global activators, MarA SoxS or Rob.

## 2. Materials and Methods

### 2.1 Bacterial Strains

All strains utilized in this study were derived from BW25113, which is the parent strain of the Keio collection [41], from which the *marA*, *soxS*, and *rob* deletion mutants were obtained. The *acrR* deletion mutant was originally obtained from the Keio collection but was reconstructed by P1 phage transduction into wild-type BW25113. The *acrR*(L34Q) mutant was constructed in our previous study [11]. The *marA*, *soxS*, and *rob* deletions were transduced into *acrR*(L34Q) using a standard PI phage transduction. The *marAsoxSrob* triple mutant was constructed by using FLP recombinase expression from the pCP20 plasmid to remove the *kan^R^* cassette from the *marA* deletion mutant, transducing the *soxS* deletion into the *marA* deletion mutant, and then repeating the above process to also delete *rob*. This process was repeated in the *acrR*(L34Q) mutant to generate the *acrR*(L34Q)*marAsoxSrob* quadruple mutant. The presence of all three deletions was confirmed using PCR. Strains CL5415 - CL5422 were constructed by transforming pBAD33, pBAD33-acrAB, pNN387, or pNN608 plasmids into electrocompetent JW5249, JW4023, JW4359. For the deletion of *acrR* past the *marbox* sequence, the Kan^R^ cassette was recombineered into BW25113 using primers 5’AGAAGCGCAAGAAACGCGCCAACACATCCTCGATGTG GCTCTACGTCTTTATGATTCCGGGGATCCGTCGACC3’and 5’CAGG AAAAATCCTGGAGTCAGATTCAGGGTTATTCGTTAGTGGCAGGATTTGTAGGCTGGAGCTGCTTCG3’All strains used in this study are listed in Table 1.

**Table 1.**
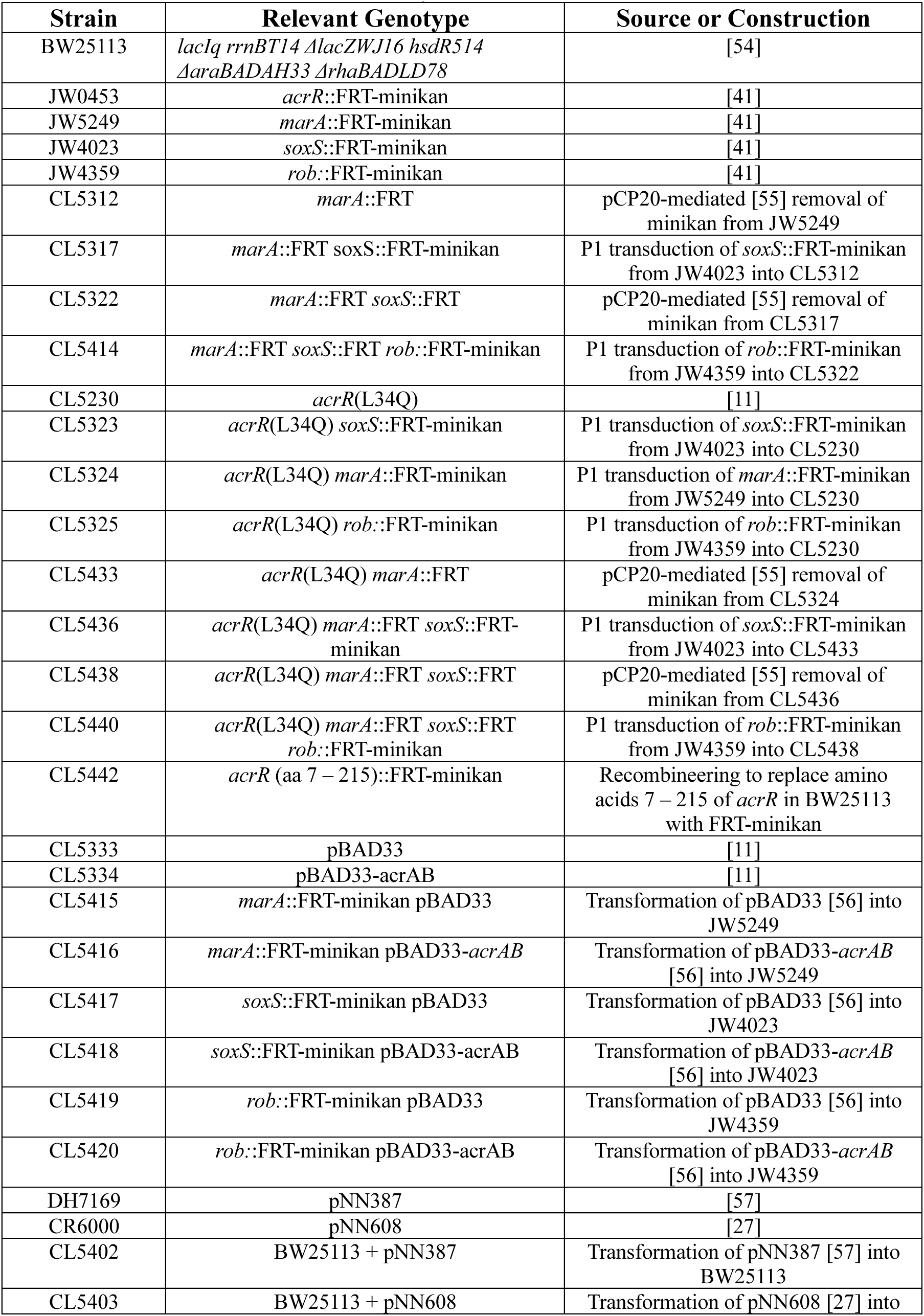
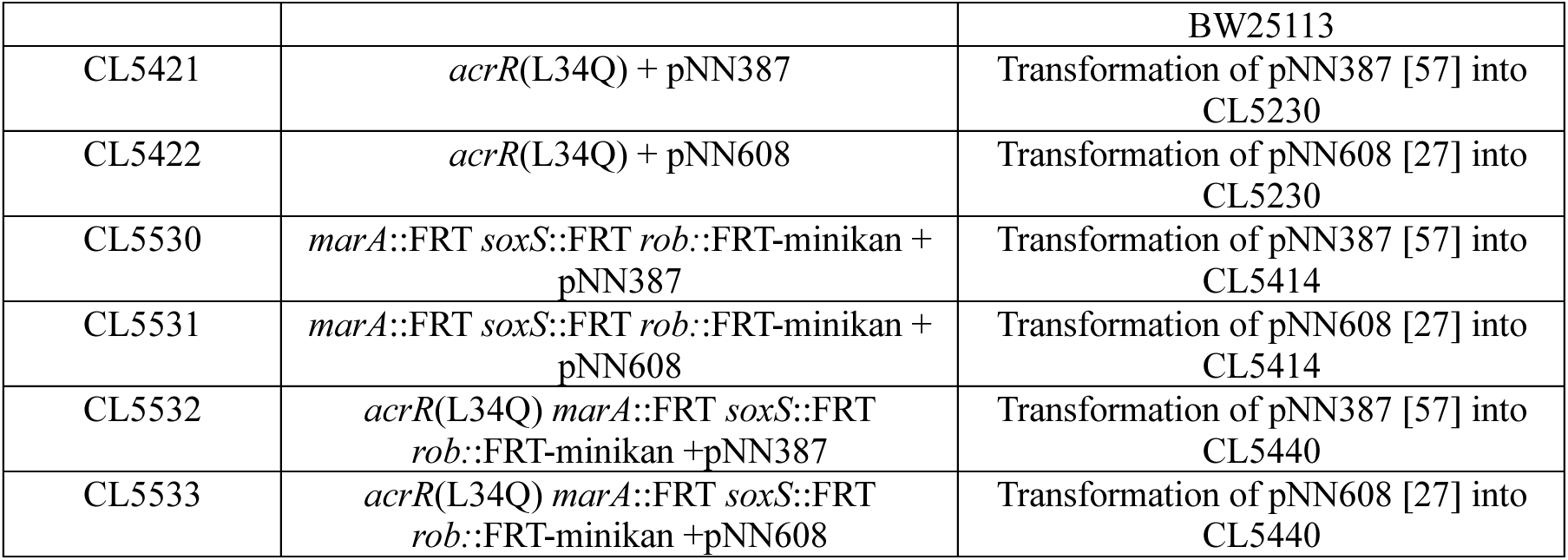
List of strains used in this study.

### 2.2 Psoralen-UVA (PUVA) survival

10-µL aliquots of 10-fold serial dilutions from overnight cultures were spotted onto LBthy plates containing 20 µg/mL 8-methoxypsoralen. Plates were then exposed to UVA irradiation at an incident dose of 6.5 J/m2/s for the indicated dose and incubated overnight at 37°C. Surviving colonies at each dose were then counted and compared to the non-exposed plates to calculate percent survival.

For overexpression of *acrAB* from expression vectors, 5 mL LB subcultures were inoculated with 50 µL of overnight cultures containing the expression plasmid, pBAD33-acrAB, or its parent vector, pBAD33, and grown in a 37°C shaking water bath to OD600 of 0.4. 1 mM L-arabinose was added to subcultures for the last 30 minutes of incubation before proceeding with survival assays as described above.

### 2.3 acrAB-lacZ expression

10-µL aliquots of 10-fold serial dilutions from overnight cultures containing pNN608 (*acrABp*-lacZ) or pNN387 (empty vector) were spotted onto LBthy plates supplemented with 120 µg/mL 5-Bromo-4-Chloro-3-Indolyl β-D-Galactopyranoside (X-Gal) either with or without 20 µg/mL 8-methoxypsoralen. Two plates each of LB X-GaL and LB X-Gal + 20 µg/mL 8-methoxypsoralen were then exposed to 3.8 kJ/m^2^ UVA radiation as described above for survival assays. Plates were then compared to unexposed plates and photographed.

## 3. Results

### 3.1 Global Regulators MarA, SoxS, and Rob are required for full resistance to psoralen-UVA

In previous work, three mutations in the transcriptional repressor *acrR* were isolated and found to confer high-level resistance to psoralen-UVA through the upregulation of *acrA* and –*B*, encoding components of the AcrAB-TolC efflux pump (Fig. 2A and [11]). However, when we deleted the entire *acrR* coding region, we found that unlike the other *acrR* mutations, no resistance was conferred (Fig. 2A). The observation argues that the loss of the AcrR repressor is insufficient to confer resistance to psoralen interstrand crosslinks and a more complex mechanism is involved in the acquisition of resistance.

**Figure 2.**
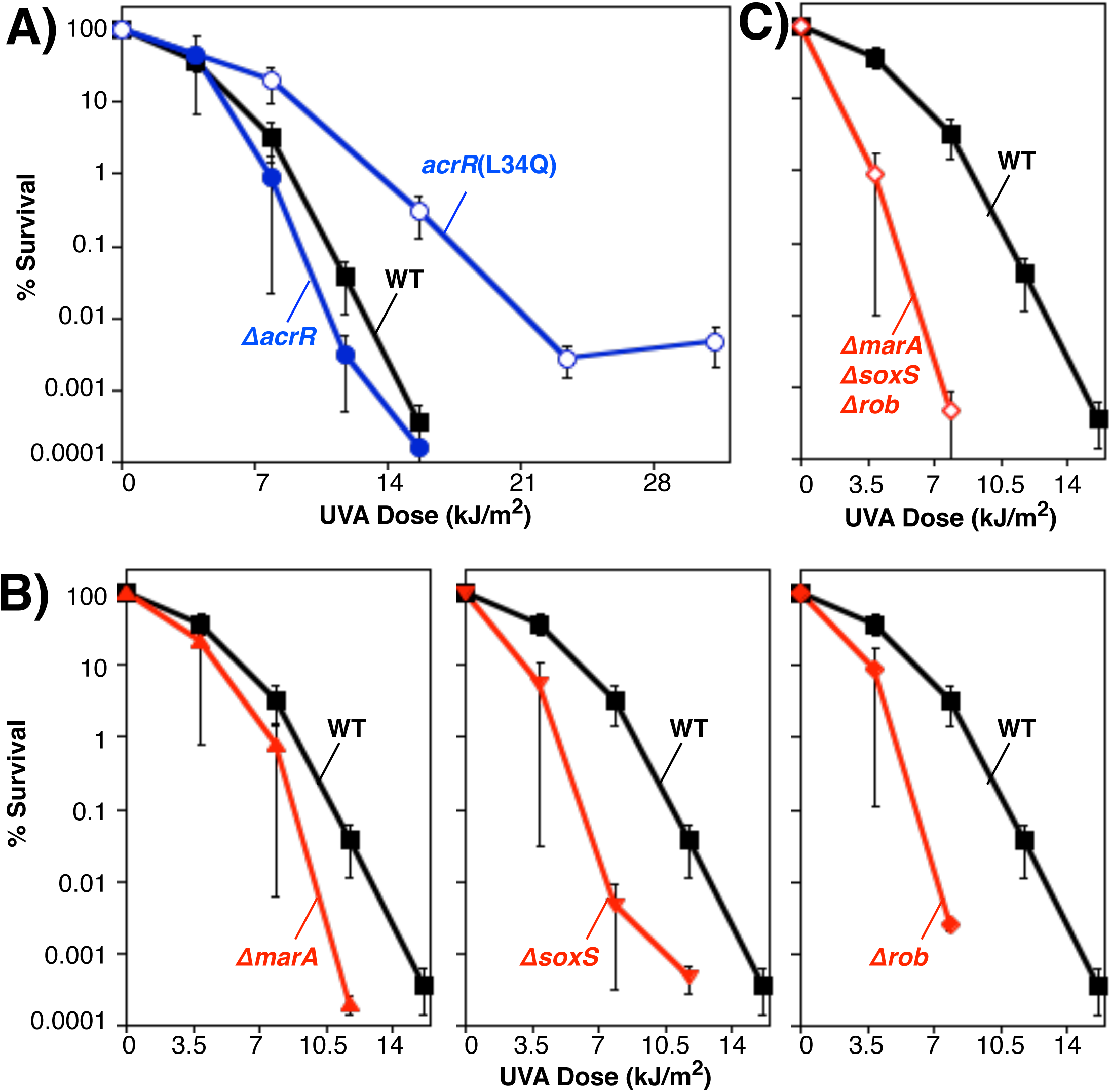
*acrR*(L34Q), but not deletion of *acrR*, confers resistance to psoralen–UVA, while deletion of *marA*, *soxS*, or *rob* renders cells hypersensitive. A) The survival of wild-type cells (filled squares); Δ*acrR* (filled circles), and *acrR*(L34Q) mutants (open circles), B) Δ*marA* (filled triangles), Δ*soxS* mutant (filled inverted triangles), Δ*rob* mutants (filled diamonds) and C) Δ*marA* Δ*soxS* Δ*rob* mutants (open diamonds) is plotted following UVA irradiation at the indicated doses in the presence of 20 µg/mL 8-methoxypsoralen. Plots represent the average of at least two independent experiments. Error bars represent the standard error of the mean. Wild type is replotted in each graph for comparison.

Common to all three of the resistance-conferring *acrR* mutants that were islaoted is that they retain the initial third of *acrR*’s coding sequence but alter or remove the latter two-thirds of the protein. The first third of the gene encodes the DNA-binding domain for the AcrR regulator. However, we also noted this region also encodes a MarA, SoxS, and Rob binding sequence, termed *marbox*, which has been reported to positively regulate the *acrA -B* operon [27, 31]. Thus, it is possible that the mutations confer psoralen resistance either through altering AcrR’s DNA binding properties or through activation of *acrAB* by MarA, SoxS, or Rob.

If the psoralen resistance is mediated through the *marbox*, then deletion of the *marA*, *soxS*, and *rob* genes would be expected to impair resistance in these strains. To test this possibility, we examined the ability of mutants deleted for these genes to survive psoralen-UVA treatment. Ten-fold serial dilutions of an overnight culture were spotted on plates containing 20 μg/mL 8-methoxypsoralen and exposed to increasing doses of UVA. Following overnight incubation at 37°C, surviving colonies were counted and compared to the unexposed plate to determine percent survival. Figure 2B shows that deletion of either *marA*, *soxS*, or *rob* renders cells more sensitive than WT to psoralen-UVA irradiation, indicating that all three of these genes are important for psoralen-UVA resistance. Notably, the contribution of each was not additive, as the absence of any single regulator resulted in hypersensitivity that was similar to the *marA soxS rob* triple mutant (Fig. 2C). Given that MarA, SoxS, and Rob all share a single *marbox* binding sequence within *acrR*, it is unexpected and remains unclear why deleting of any one of these three proteins renders cells hypersensitive. However, the observation indicates that all three proteins are required to maintain resistance to psoralen-UVA interstrand crosslinks, despite sharing a single DNA binding sequence.

### 3.2 MarA, SoxS, and Rob contribute to psoralen-UVA resistance primarily through upregulation of acrAB

MarA, SoxS, and Rob upregulate expression of approximately 50 genes in response to various cellular stresses [31]. Thus, although the results of Fig. 2 indicate that MarA, SoxS, and Rob are required for full resistance to psoralen, they do not establish if this contribution can be attributed directly to the upregulation of *acrAB* or if resistance is conferred by other *marbox*-regulated genes. To test this, we used an arabinose-inducible *acrAB* plasmid to overexpress *acrAB* in the *marA*, *soxS*, *rob* deletion mutants, which would result in upregulation of *acrAB*, but not any other *marbox*-regulated genes. Actively growing cultures containing the plasmid were incubated with arabinose for 30 minutes to induce *acrAB* expression prior to psoralen-UVA treatment. Figure 3 shows that plasmids containing the *acrAB* sequence increase resistance in *marA*, *soxS*, and *rob* mutants to near wild-type levels. By contrast, mutants containing an identical plasmid lacking the *acrAB* sequence remain hypersensitive to psoralen-UVA treatment. The results indicate that MarA, SoxS, and Rob contribute to psoralen-UVA resistance primarily through upregulation of *acrAB* expression and that the idea that loss of this upregulation in the *acrR* deletion mutant could be responsible for its inability to confer resistance.

**Figure 3.**
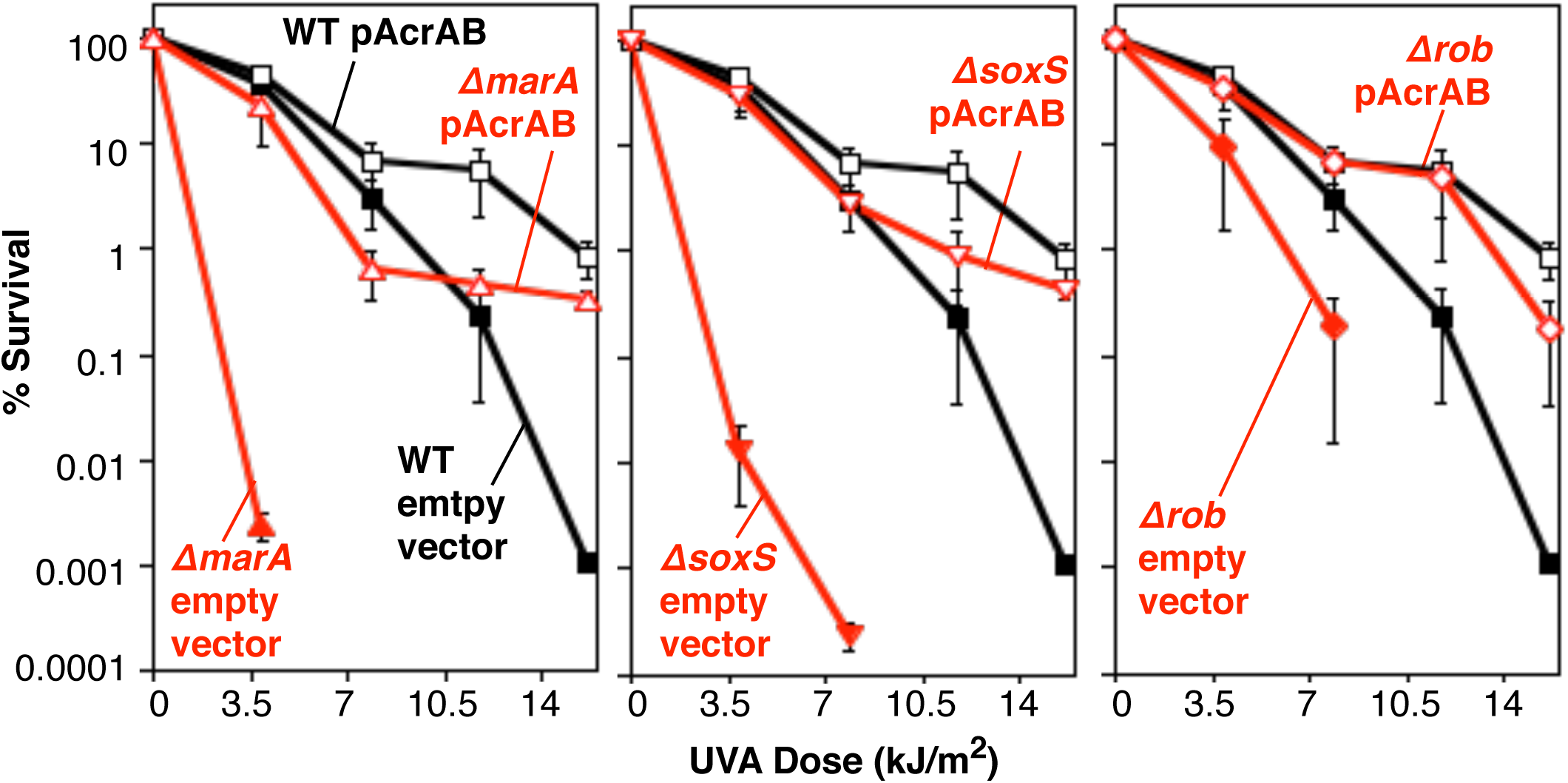
Overexpression of AcrAB alone is sufficient to restore psoralen-UVA resistance in Δ*marA*, Δ*soxS*, and Δ*rob* mutants. The survival of wild type (squares), Δ*marA* (triangles), Δ*soxS* (inverted triangles), and Δ*rob* (diamonds) containing either an empty pBAD33 expression vector (filled symbols) or an AcrAB expression vector (open symbols) is plotted following UVA irradiation at the indicated doses in the presence of 20 µg/mL 8-methoxypsoralen. Plots represent the average of at least two independent experiments. Error bars represent the standard error of the mean. Wild type is replotted in each graph for comparison.

### 3.3 MarA, SoxS, and Rob activation and AcrR de-repression contribute additively to psoralen interstrand crosslink resistance

MarA, SoxS, and Rob-mediated upregulation of *acrAB* is required for full resistance to psoralen-UVA (Fig. 2 and 3). Since the highly resistant *acrR*(L34Q) mutant retains the *marbox* sequence, it is possible that the high level of resistance requires activation by MarA, SoxS, or Rob. If true, we would expect that deletion of *marA*, *soxS*, or *rob* would significantly reduce psoralen-UVA resistance in the *acrR*(L34Q) strain. As shown in Fig. 4, *acrR*(L34Q) mutants remained resistant to psoralen-UVA, when either *marA*, *soxS*, or *rob* was deleted. However, the loss of all three genes reduced the resistance of *acrR*(L34Q) mutants to levels similar to wild-type cells and the *acrR* deletion mutant. Taken together with the previous observations, the results support the idea that both de-repression by AcrR and activation by MarA, SoxS, or Rob are required to achieve resistance to psoralen interstrand crosslinks.

**Figure 4.**
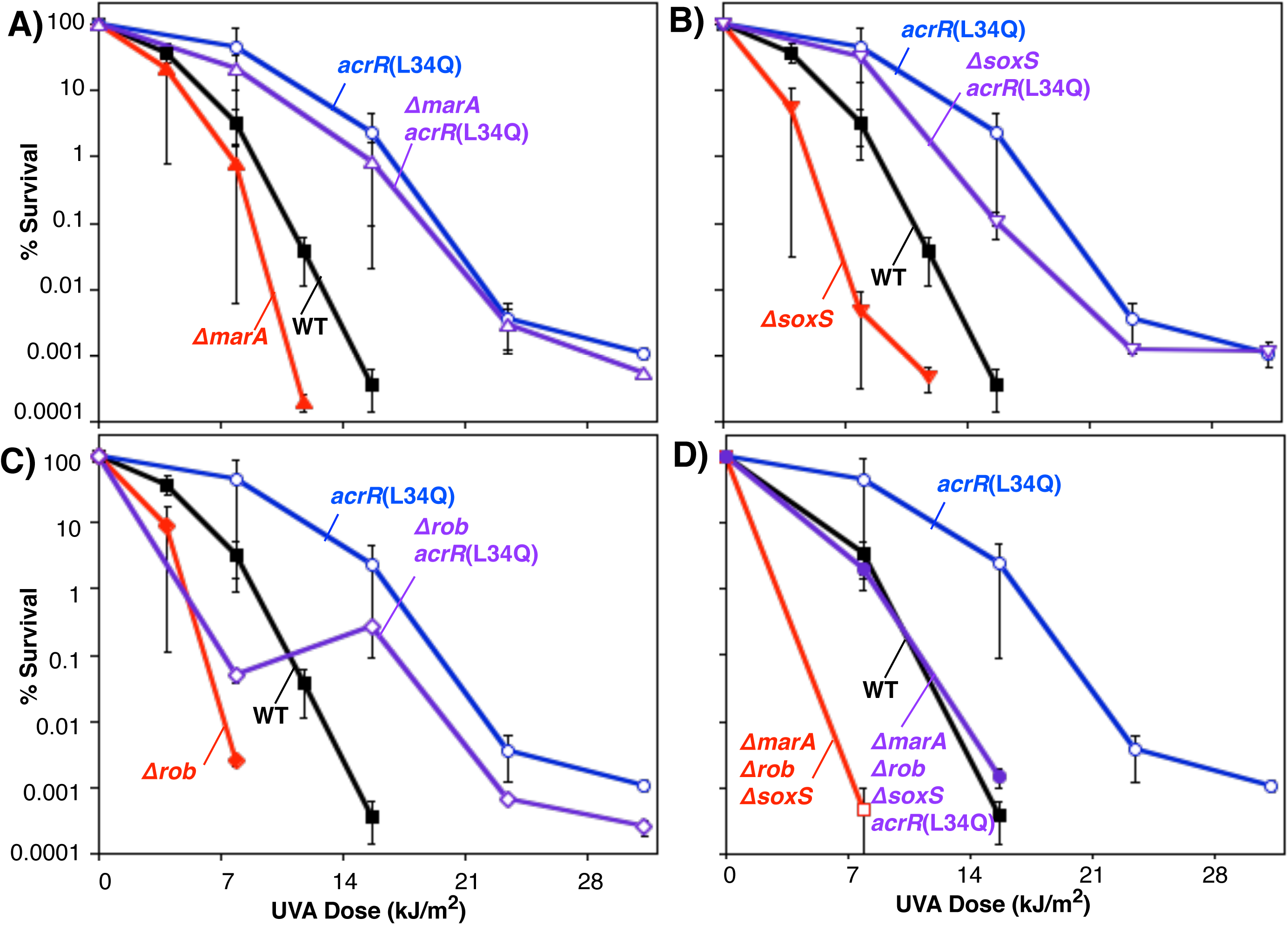
*acrR*(L34Q) psoralen-UVA resistance requires MarA, SoxS, and Rob activation. The survival of wild-type cells (filled squares), *acrR*(L34Q) (open circles), Δ*marA* (closed triangles), *acrR*(L34Q) Δ*marA* (open triangles); (B) Δ*soxS* (filled inverted triangles), *acrR*(L34Q) Δ*soxS* (open inverted triangles); (C) Δ*rob* (filled diamonds), and *acrR*(L34Q) Δ*rob* (open diamonds); (D) Δ*marA* Δ*soxS* Δ*rob* mutants (open squares), *acrR*(L34Q) Δ*marA*Δs*oxS*Δ*rob* (filled circles) in the presence of 20 µg/mL 8-methoxypsoralen at the indicated UVA doses is plotted. Plots represent the average of at least two independent experiments. Error bars represent the standard error of the mean. Wild type is replotted in each graph for comparison.

To confirm these requirements directly, we used recombineering to generate a complete deletion of the *acrR* coding sequence with the exception of the first 21 nucleotides encoding the *marbox* sequence. Figure 5 shows that the *marbox* sequence alone is sufficient to restore full resistance to *acrR* deletion mutants, mimicking the resistance seen in the *acrR*(L34Q) mutant.

**Figure 5.**
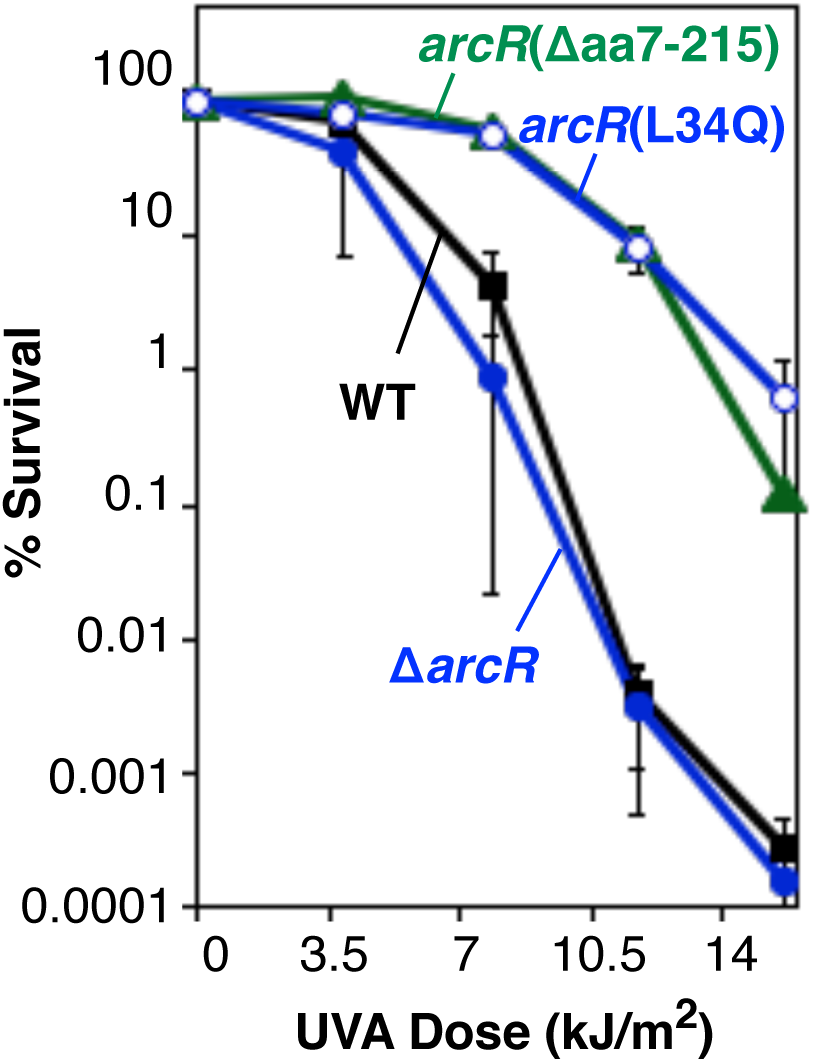
The *marbox* sequence is necessary and sufficient to induce resistance to psoralen interstrand crosslinks in the absence of the AcrR repressor. The survival of wild-type cells (filled squares), Δ*acrR* (filled circles), *acrR*(L34Q) (open circles), and Δ*acrR* (aa7-215) (filled triangles) in the presence of 20 µg/mL 8-methoxypsoralen at the indicated UVA doses is plotted. Plots represent the average of at least two independent experiments. Error bars represent the standard error of the mean.

### 3.4 acrAB expression is not induced by psoralen, UVA, or psoralen-UVA irradiation

The results above demonstrate that the AcrR transcriptional regulator and activation by either MarA SoxS or Rob are required toupregulate *acrAB* and confer to crosslink resistance. However, how *acrAB* is regulated in wild-type cells during challenge with psoralen-UVA is unknown. Previous studies have shown that exposure to others stressors and agents-including ethidium bromide, cadaverine, ethanol, or high osmolarity, can induce expression of *acrAB* to increase resistance [27, 42]. To examine if *acrAB* expression is responsive to psoralen-UVA treatments, we used a plasmid that contained the *acrAB* promotor region fused to *lacZ*. The cloned promoter region contains both the AcrR binding site as well as the first 102 nucleotides of *acrR* coding sequence which contains the *marbox* binding site. To test if psoralen, UVA irradiation, or the presence of interstrand crosslinks can serve to induce *acrAB* expression, cultures containing the plasmid were spotted in 10-µL serial dilutions on X-Gal plates that were left untreated or exposed to either psoralen, UVA, or psoralen-UVA. As shown in Figure 6A, in the presence of the *acrABp-lacZ* reporter, the parental strain detectably expressed the *acrAB* genes as indicated by the partially blue colonies, relative to the control plasmid. As controls, we also examined the *acrR*(L34Q) resistant mutant and the sensitive *marA soxS rob* deletion mutant. As expected, *acrAB* expression was elevated in *acrR*(L34Q) mutant as indicated by the intensely blue colonies, correlating with the increased expression of *acrAB* and resistance in this strain. Similarly, colonies were noticeably less blue in the *marA soxS rob* deletion background which correlates with reduced *acrAB* expression and hypersensitivity.

**Figure 6.**
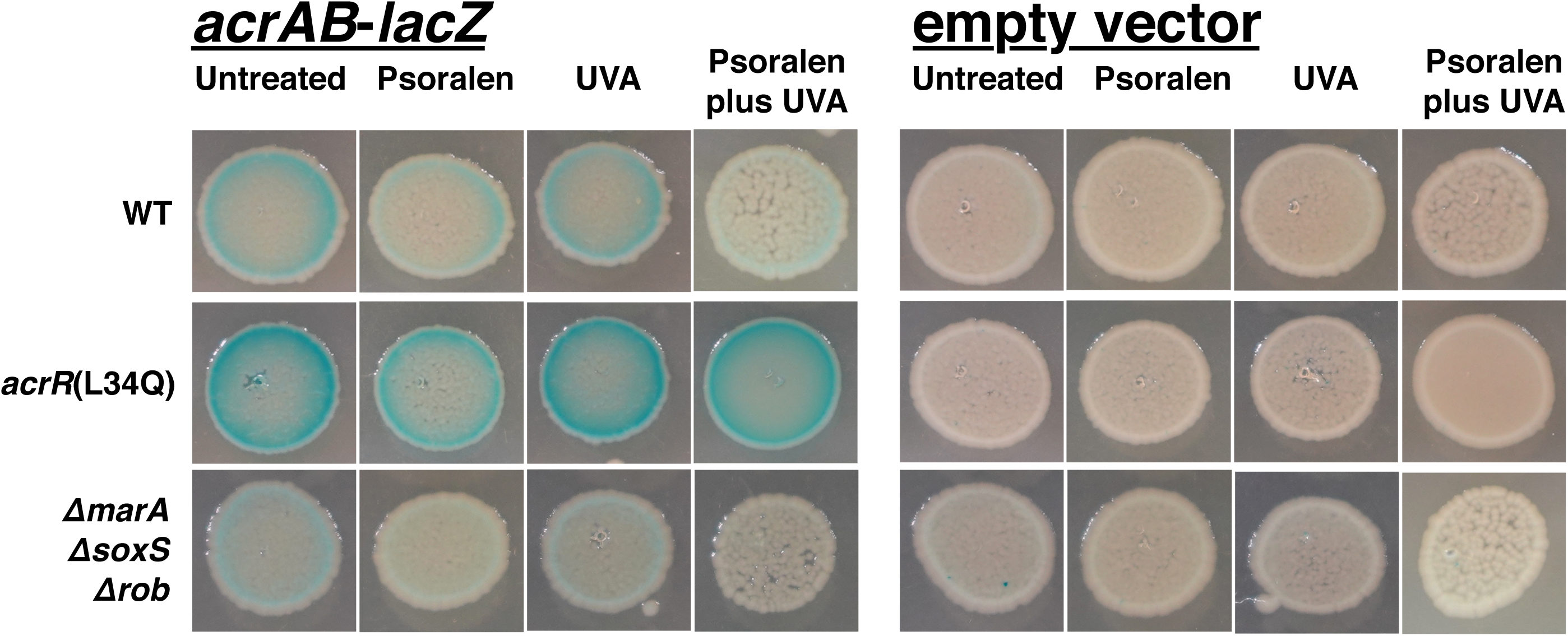
*acrAB* expression is not upregulated by psoralen, UVA, or psoralen-UVA. 10 µL spots of 10^4^ cells of wild type, *acrR*(L34Q), or Δ*marA* Δ*soxS* Δ*rob*, mutants containing a LacZ reporter plasmid fused with the *acrAB* promoter region (p-*acrAB*-*lacZ*) or no promoter region (empty vector) were plated on LB plates containing X-Gal. Plates contained 20 µg/mL 8-methoxypsoralen and were UVA irradiated with 3.8 kJ/m2 as indicated. LacZ expression from the plasmids is indicated by blue color in colonies.

Notably however, expression did not increase in the presence of either psoralen, UVA, or psoralen plus UVA treatments. The results imply that psoralen, UVA, or the combination do not generate substrates that activate *acrAB* and suggest these regulators are not normally responsive to this challenge, in the absence of mutation.

## 4. Discussion

The results demonstrate that all three of the related global regulators MarA, SoxS, and Rob have a significant role in psoralen-UVA resistance. However, regulation by these activators was found to be complex. Deletion of any single global effector gene in wild-type cells had a similar impact on psoralen-UVA resistance as deleting all three genes. This result is unexpected for several reasons. First, although *rob* is expressed constitutively, *marA* and *soxS* are expressed at relatively low levels until a specific stressor induces their expression [31, 32, 34, 35, 39, 43]. Additionally, the activity of Rob has been shown to be responsive to its recognition of various substrates and its release from sites of sequestration in the cell [40, 44]. Yet despite these regulators responding to different stressors, Fig. 2 demonstrates that no single effector is responsible for initiating the stress response to psoralen-UVA irradiation. Second, given the high level of homology between MarA, SoxS, and Rob, and their ability to bind the same *marbox* sites across the genome, one might expect that loss of one regulator could be offset by the presence of the other two [31, 32, 34, 35, 39, 45]. Yet this is not observed in wild-type cells. On the other hand, if no redundancy existed, one might expect that deleting *marA*, *soxS*, and *rob* would have an additive effect on psoralen-UVA sensitivity. This is also not observed (Fig. 2). Thus, the apparent any-and-all requirement for MarA, SoxS, and Rob could suggest that crosstalk between these activators is particularly important in psoralen-UVA resistance. Alternatively, it is possible that the sensitivity of this assay to distinguish phenotypic differences decreases as the limits of detectability are approached.

Irrespective of the crosstalk, the AcrAB-TolC transporter appears to be the causal target for MarA, SoxS, and Rob generating psoralen crosslink resistance, since the hypersensitive phenotype of these mutants can be suppressed by overexpression of AcrAB alone and does not require any of the other approximately 50 genes under their regulation [31–40]and Fig 3).

The results may also suggest a more complex mechanism of regulation by AcrR than that of a simple repressor. The *acrR*(L34Q) point mutation is resistant to psoralen-UVA treatment and retains the *marbox* activation sequence, yet a deletion of the *acrR* open reading frame that deletes the *marbox* activation sequence renders cells sensitive. If AcrR acts as a basic repressor, then the simplest model would be that *acrR*(L34Q) is a null mutation, and that upregulating *acrAB* expression enough to provide full resistance requires both removal of the AcrR repressor and activation by MarA, SoxS, and Rob. The finding that removal of all three proteins renders the *acrR*(L34Q) mutant similar in resistance to the *acrR* deletion supports this model (Figure 4D). In contrast to what was seen in wild-type cells (Fig 2), deletion of *marA*, *soxS*, or *rob* individually was insufficient to noticeably reduce resistance in *acrR*(L34Q). As mentioned above, one possible explanation would be that the sensitivity of the survival assays used in this study are insufficient to distinguish small differences between hypersensitive strains, as their numbers approach the lower end of detectability. If true, increasing the background level of resistance through de-repression of *acrAB* expression as in Figure 4 could make the differences between individual *marA*, *soxS*, and *rob* deletion mutants and the *marAsoxSrob* triple mutant more detectable.

The most prominent model for AcrR repressor function is that it releases upon binding a recognized substrate [27–29, 46, 47]. These initial studies used both LacZ reporter constructs and gel-shift binding assays to provide strong evidence that AcrR can repress expression of *acrAB* when bound to its promoter [27]. However, other aspects of this study also suggested more complexity in its function. Transcription of the *acrR* repressor was also induced by the same stressors that upregulated *acrAB* transporter expression, and *acrAB* upregulation during ethanol stress or growth phase occurred independently of MarA and SoxS, similar to what we observe in the presence of psoralen [27]. The repressor model of AcrR activity is based on its similarity to other TetR family transcriptional regulators. It proposes that upon ligand binding, AcrR releases from DNA to allow transcription. However the AcrAB-TolC transporter is active on a wide range of structurally divergent substrates [42, 47], making it unclear how the protein could effectively recognize this diverse range of toxic substrates. The few substrates which have been examined and shown to promote AcrR release from oligos *in vitro* have been DNA intercalators [28, 29, 46] which makes it difficult to determine if release is due to ligand binding or changes to the DNA structure of the oligos used. Further, studies looking for upregulation of AcrAB following treatments with known substrates of the pump have seen modest to no effect [42, 47]. Thus, the mechanism of regulation and natural substrate for induction of the efflux pump remains unclear.

This ambiguity of the regulatory mechanism also holds true for resistance to psoralen interstrand crosslinks. Although psoralen is clearly a substrate for the efflux pump [11], AcrR does not appear to be naturally responsive to this stress, as neither psoralen, UVA, nor crosslink induction upregulates *acrAB* expression. This perhaps makes sense given that the cell must acquire mutations that disrupt AcrR repression of the efflux pump to achieve resistance. Importantly, we show here that full resistance is only achieved when *acrR* null mutations preserve the *marbox* sequence (Figure 5), as in *acrR*(L34Q). This is particularly relevant to the emergence of multi-drug resistance in Gram-negative bacteria, as mutations in *acrR* are known to drive multi-drug resistance and are commonly found in clinical isolates [48–52]. Additionally, the original studies that characterized AcrR used insertion mutants that disrupted the protein after the *marbox*, while later studies used the complete deletion mutant of *acrR* from the Keio collection [26, 27, 41, 42, 47]. Our results demonstrate that it will be important in the future to consider the impact of *acrR* mutations on the *marbox* when assessing their phenotypic effects. Finally, it is also notable that the resistance to psoralen interstrand crosslinks is achieved by preventing this drug from forming this lesion. No mutations upregulating repair pathways or proteins were observed in the initial screen, consistent with previous studies that found cells lack effective repair mechanisms for this form of damage [6–8, 53].

## Ethical approval

Not required.

## Data Availability

All data is contained within the manuscript. Data, strains, and plasmids used in this work are available upon request to the corresponding author.

